# Machine-learning for cluster analysis of localization microscopy data

**DOI:** 10.1101/505719

**Authors:** David J. Williamson, Garth L. Burn, Juliette Griffie, Daniel M. Davis, Dylan M. Owen

**Affiliations:** Randall Centre for Cell and Molecular Biophysics, King’s College London, London UK; Max Planck Institute for Infection Biology, Berlin Germany; Laboratory of Experimental Biophysics, EPFL, Lausanne Switzerland; Manchester Collaborative Centre for Inflammation Research, University of Manchester, Manchester UK

## Abstract

Quantifying the clustering of points within single-molecule localization microscopy data is useful to understanding the spatial relationships of the molecules in the underlying sample. The conversion of point pattern data into a meaningful description of clustering is difficult, especially for biologically derived data, as the definitions of clustering are often subjective or simplistic. Many existing computational approaches are also limited in their ability to process large-scale data-sets or to deal effectively with inhomogeneities in clustering. Here we have developed a supervised machine-learning approach to cluster analysis which is fast and accurate. Trained on a variety of simulated clustered data, the network can then classify all points from a typical localization microscopy data-set (several million points from the entire field of view) as being either clustered or not-clustered, with the potential to include additional classifiers to describe different types of clusters. Clustered points can then be further refined into like-clusters for the measurement of cluster area, shape, and point-density. We demonstrate the performance on simulated data and experimental data of the kinase Csk and the adaptor PAG in both naive and pre-stimulated primary human T cell synapses.

## Introduction

Single-molecule localization microscopy (SMLM) brings many challenges, from specimen preparation to the acquisition and reconstruction of the data into a high fidelity super-resolution image. Once a super-resolved SMLM image has been obtained there is the further challenge of how to effectively analyse such data. Whereas images from wide-field or confocal laser scanning microscopes are an array of picture element intensity values, the data from SMLM is fundamentally a list of coordinates, each representing the spatial location of a point-emitter within the original specimen. This list of points can be plotted and rasterized for examination with conventional image analysis tools but an ideal analysis method would operate on the original coordinate data without requiring its transformation to another format. As SMLM data is a distribution of points within the imaging field it can be interrogated using spatial point pattern analyses to reveal the spatial relationships between the points and also higher-scale relationships between clusters of points, or between points from a different target or imaging channel. Such techniques include Ripley’s *K* Function (*Ripley 1977*), it’s related transformation as Besag’s *L* Function (Besag 1977), and a local generalisation as Getis & Franklin’s local point pattern analysis (Getis and Franklin 1987), as well as the radial distribution (or pair correlation) function (Howroyd, Stoyan, and Stoyan 1996), and DBSCAN (“A Density-Based Algorithm for Discovering Clusters in Large Spatial Databases with Noise” 1996). Common among these approaches is to operate on each point over a set of distances (or a fixed distance) and compare the number of other points found to the number of points expected to be found. This requires judicious selection of analysis parameters which can lead to a sub-optimal interpretation of the data, for example when points are clustered at a different spatial scale to the one used for assessment. The analysis can be confounded if points are inhomogeneously clustered, for example if a large, dense concentration of points (such as a position registration bead) falls within the region under interrogation or there are multiple types of clusters present. These situations are common in the case of complex data from biological specimens and the selection of appropriate analysis parameters is a time-consuming and qualitative undertaking. In addition, these approaches often require a threshold to distinguish clustered from non-clustered points and it is often the case that the appropriate threshold for one image is entirely unsuitable for the next, even within images from the same experimental condition. Furthermore many of these methods are only practical to use for small number of points, with larger datasets saturating or exceeding the available computing resources. These problems are usually avoided by reducing the input data, for example by only considering a subset of the total points within a small region of interest. This approach can result in missing important broader cellular contexts, or introducing user-bias in the placement or selection of regions-of-interest. The image can be broken down into multiple adjacent (or overlapping) regions, analysed per-region, and then reconstituted but this can often exacerbate the problem of selecting the analysis parameters which are the most appropriate for each region.

In spite of the numerous mathematical approaches to identify clusters within a field of points, such a task is often very easily undertaken manually; human perception is naturally adept at identifying clustered points and cluster boundaries within images. The limitation with a manual approach is simply one of patience and mental exhaustion as there can be thousands to millions of points in a single image, which may group into hundreds to thousands of clusters. Machine learning is a computational and statistical approach to extract meaningful information from complex data where a fully descriptive model is not otherwise available. One of the tools used in machine learning is that of neural networks which were so named as they were inspired by biological neurons which are self-contained units capable of accepting one or more input signals which are then integrated and processed internally to generate one or more output signals. These output signals can then be used as input for a second set of neurons, eventually resulting in a system which can generate output as an abstract representation of the original input. Machine learning has developed rapidly within the field of artificial intelligence where it has been predominantly employed in the service of problems such as facial recognition, autonomous vehicle navigation, natural language interpretation and synthesis, speech recognition.

Machine learning, when applied to microscopy images often uses convolutional neural networks to examine raster-based images. Such networks were originally developed and refined to solve computer vision problems and have proven very useful in the identification and extraction of abstract features from images. For microscopy, they are employed to good effect in finding shapes and structures within optical(Hay and Parthasarathy 2018) and electron microscopy(Roels et al. 2017) data. Machine learning has also been used to enhance the spatial resolution of images from conventional microscopes(Rivenson et al. 2017) and also to derive super-resolved images from partial sets of localization microscopy data (Ouyang et al. 2018). However, as localization microscopy data is not immediately compatible with convolutional neural networks a different approach is required. Machine learning is still a useful approach as the goal of segmenting a pointillist data-set into clusters is ultimately one of classification in which a point is labelled as being from a cluster or not from a cluster.

Here we employ machine learning to solve realistic cluster identification problems in SMLM data-sets by training a neural network to extract features from nearest-neighbour distance-derived data. A set of software modules is presented to prepare raw data, train new models, evaluate data with trained modules, form cluster shapes around clustered points identified by models, and extract descriptive data from the clusters and clustered points. A quantitative comparison to other methods demonstrates the accuracy of our approach, with particular benefits of being able to access arbitrary regions of interest from very large data-sets and with a comparably fast processing time.

## Methods

For a machine learning model to be trained, it must be given input data to work with. For a point within a spatial point-pattern a sequence can be constructed of the distances from the point to its neighbouring points, from the nearest neighbour (NN_1_; the zero^th^ neighbour NN_0_ is taken to mean the point itself) to the n^th^ nearest neighbour (NN_n_), and where n is less than the total number of points in the field. For an arbitrary point within a field of randomly distributed points, there ought to be a different pattern to these distance values compared to a point within a locally dense region (i.e. a cluster). For clustered points, the distances to the nearest neighbouring points should be relatively consistent for those nearby points which are also within the cluster, but this pattern should change once further neighbours, found outside of the cluster, are accessed (Fig. 1a-d). Similarly, points on the edges of clusters will find their near-neighbours lie within that cluster and further neighbours outside of the cluster, depending on the size of the cluster and the density of non-clustered points. The monotonic sequence of near-neighbour distances for a single point, or the difference in distances between consecutive near-neighbours, can be used as input for a machine learning model.

### Workflow

Python code was written to generate data for training and to prepare and carry data through the analysis workflow from the initial preparation stages to the final collection of clustering statistics. Models were constructed from layers using Keras, an open-source machine-learning framework for Python (Chollet and others 2015). Keras makes it straightforward to construct machine learning model configurations by assembling layers, with the output of each layer being passed as input to the next. The workflow and the Python scripts used at different stages is shown in Fig. 1e. In the absence of a trained model, data with known clustering characteristics are simulated. These data are prepared for training by measuring the distances for each point to its nearest number of specified neighbours. Next, a model is specified in Keras; three examples of accurate models are described here but it is possible for the user to configure any model, as required. Each model is assigned a unique six character identity for ease of reference. Here we describe several models with different configurations utilising different near-neighbour distance-derived data as input. Model XPILJZ (Fig. 1f) consists of an input layer for 100 near-neighbour values followed by two fully connected layers and an output layer. Model 07VEJJ also requires 100 near neighbour input values but uses a more complicated configuration with a 1-dimensional convolution layer, maximum pooling layer, a dropout layer for regularization, two long short-term memory (LSTM) layers, followed by more dropout, max-pooling, and fully connected layers before the output layer which produces a single score value. It was expected that the LSTM layers might increase the models’ classification accuracy because the sequential nature of the input data are ideal input for LSTM networks (Hochreiter and Schmidhuber 1997). An LSTM is a type of recurrent neural network (RNN) formed from a chain of network units which allow information to persist (as ‘memory’) and allow the network to learn long-term dependencies from the input sequence. Similarly, Model 87B144 (Fig. 1g) employs a similar configuration as 07VEJJ but requires 1000 near-neighbour distance-derived values as its input.

**Figure 1.**
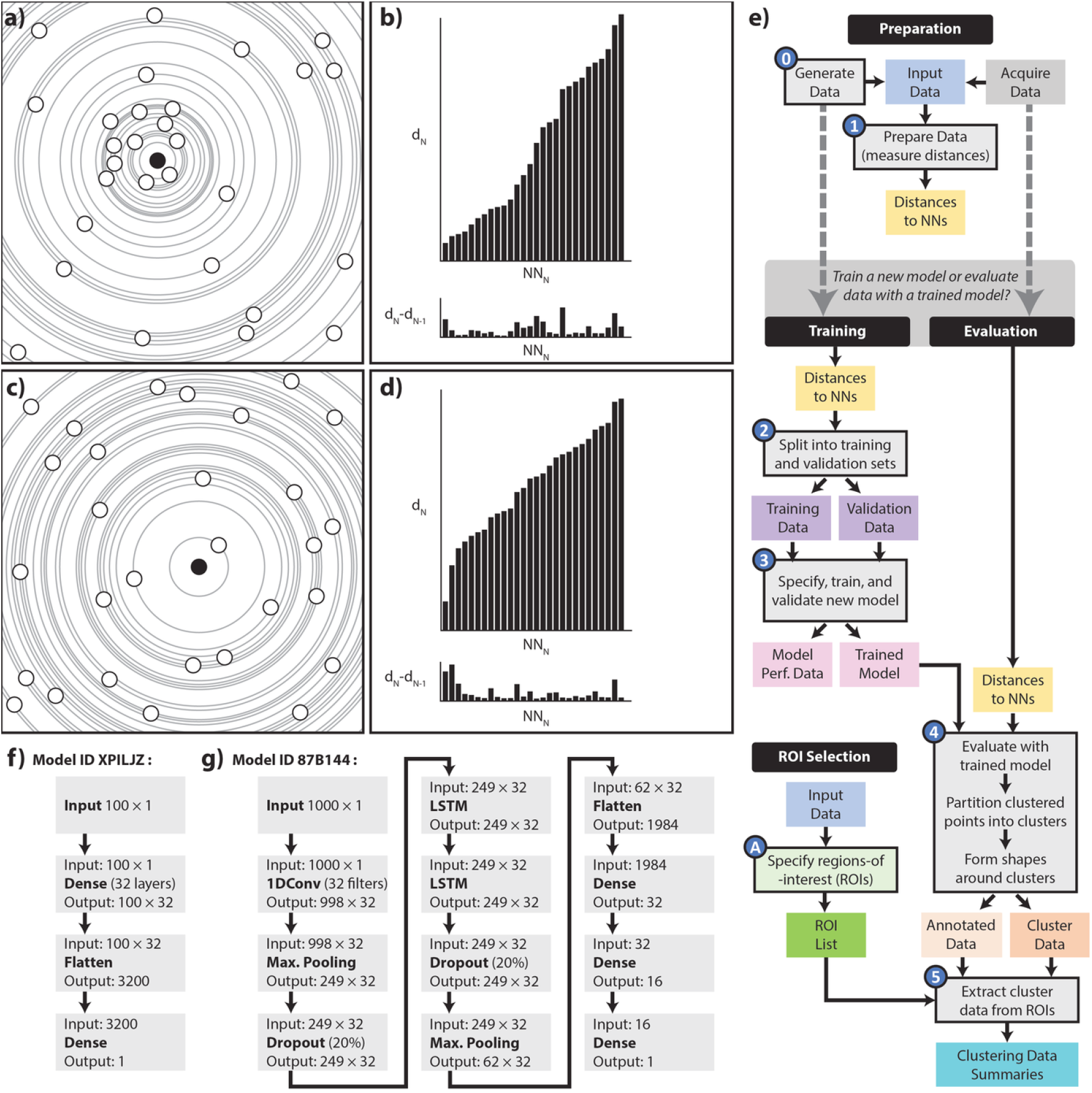
a) A collection points in a clustered configuration with a reference point highlighted in solid black, b) The distances from the reference point to its consecutive neighbours (top) and the difference between consecutive distance (bottom). (c, d) Points in a non-clustered arrangement and distances for a single highlighted point. The features within a point’s nearby-neighbour distances and distance-differences can indicate if that point is in a clustered or non-clustered pattern. e) Workflow; coloured shapes indicate input or output files. Shapes with black border are stages handled by software described here. f, g) show Keras configurations for models XPILJZ and 87B144.

### Provision of training data for supervised learning

Supervised learning approaches requires a substantial amount of labelled training data and to satisfy this requirement in the absence of manually annotate clustered data we instead simulated such data. Training data were drawn from a pool of points extracted from input data-sets, each simulated according to a different clustering scenario, e.g. ‘100 points per μm^2^, 50% of points in clusters, and 10 points per cluster’. These simulated data-sets were generated to resemble the expected format of input data from the user, specifically a character-delimited text file such as might be generated by many popular SMLM reconstruction software. Each data-set had a defined clustering of it constituent points and this clustering scenario was drawn from the combination of a range of clustering parameters describing the distribution of points within clusters and within the ‘image’ overall. At a minimum each data-set file contained a pair of x- and y-coordinates, arranged in columns, for each point. Simulated data included additional values indicating the unique cluster ID to which that point belonged, or zero to indicate a non-clustered point. The points were contained within a two dimensional ‘field of view’ comparable to that of an SMLM instrument (40 × 40 μm). The distribution of points was further restricted within a ‘cell-like’ shape, mimicking a T cell synapse formed against a flat surface and imaged by total internal reflection fluorescence (TIRF) microscopy. This was done to represent the type of gross morphology seen in a typical cell-synapse SMLM image, such as membrane protrusions, uneven edges, and other constrained geometries. The presence of these features would supply the model with training data from a context not too dissimilar to the real data that will be evaluated with the trained model.

Within the cell-like shape, points were distributed according to a specified clustering scenario. For scenarios containing clustered points, the total number of clusters was determined and a set of coordinates, each representing the seed-point for a cluster, was randomly distributed within the image field. Seed-points which fell outside of the cell-boundary shape or which were inside but located too close to either the cell outline (i.e. within the designated maximum cluster radius) or to another cluster seed (i.e. within the twice the maximum cluster radius) were randomly repositioned until all the required seed points were satisfactorily located. Clustered points were then distributed around each cluster-seed according to that scenario’s specifications. The distribution of points was uniform about the seed to generate disc-shaped (hard-edged) clusters although the potential exists to distribute point about the seed in any manner desired, for example a constant central density with a truncated Gaussian edge may better represent actual arrangement of clustered proteins in a real biological context. The number of non-clustered points was determined and these points were uniformly distributed within the cell-shape but not too close to one of the cluster seeds so as to preserve the clustering properties of the final set of points. Finally a delimited text file was saved containing the coordinates of all points, the cluster label for each point (0 for non-clustered and 1 for clustered), and a cluster ID value indicating to which specific cluster a point belonged. This process was repeated for each replicate of the cluster scenario and for all cluster scenarios; no two generated images shared either the same ‘cell outline’ or the same x-y arrangement of cluster seeds or points. Simulated data were also generated for the validation and performance testing of trained models.

**Figure 2.**
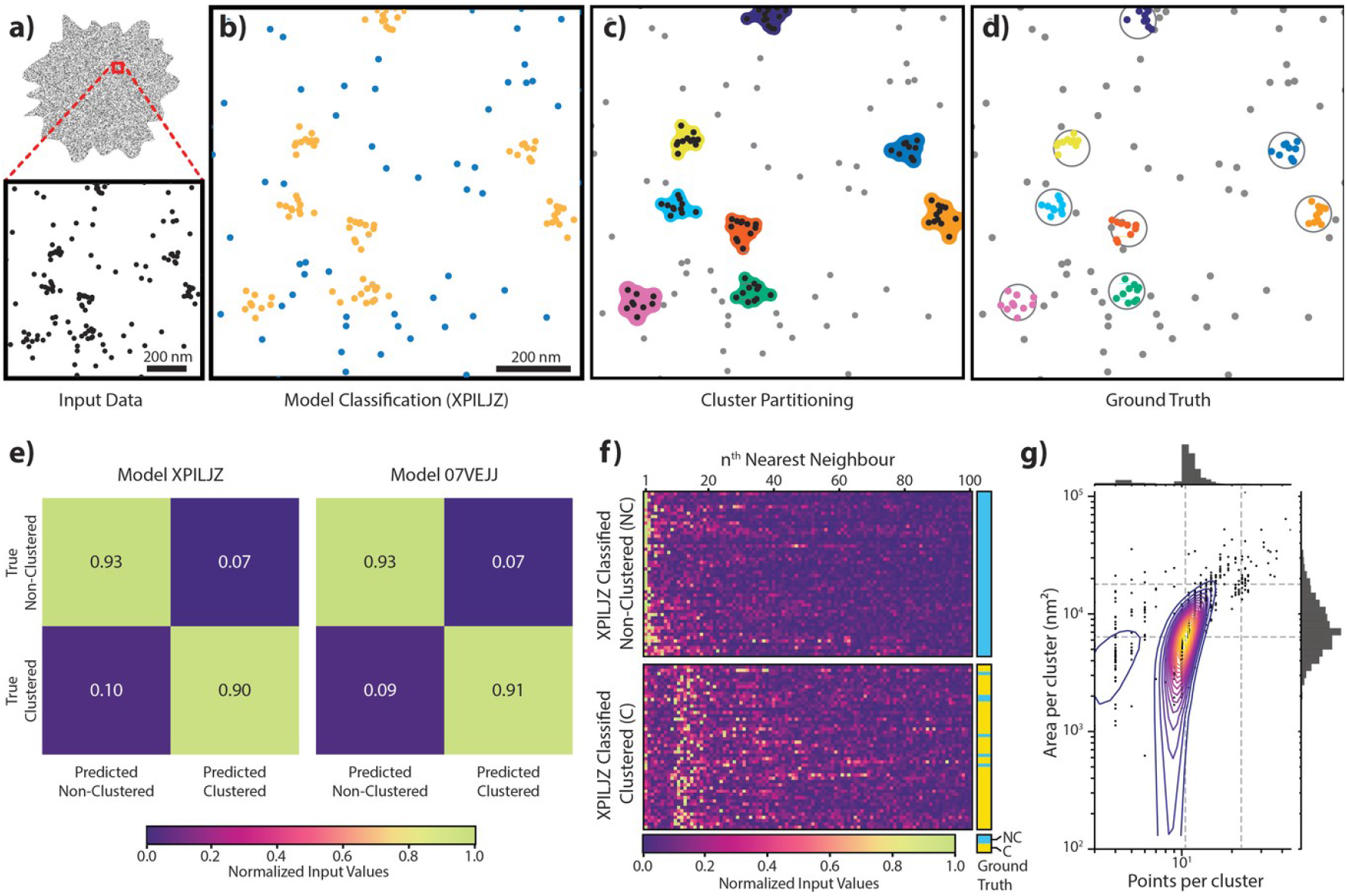
Validation of two models using simulated data. (a) Input data are the point coordinates, which are (b) evaluated by a model and classified as either clustered (gold coloured spots) or non-clustered (blue spots). This extra information is then used to partition the clustered points into spatially similar clusters, around which a cluster shape can be formed (c). Comparing the output clustering assignment to the original (known) clustering assignment (d) allows the accuracy of the process to be determined (e). Confusion matrices for two models generated using different configurations show good validation accuracy. (f) A random sample of the input data (normalized distance-differences to 100 near neighbours) for the simulated cell shown in the top row, separated into those points which were classified as nonclustered (top) or clustered (bottom); the true label for each point is shown on on the right (blue: non-clustered, yellow: clustered). (g) output data faithfully describes the original clustering configuration, here the clusters contained 10 points within a maximum radius of 40 nm.

### Preparation of input data for training and evaluation

Large data-sets can become a problem for analysis methods which attempt to operate upon all points at the same time, exhausting the available computational resources. Strategies to effectively manage larger data-sets can include acquiring more powerful computing resources or breaking a large data-set into manageable chunks and recombining the results after processing is complete. Some approaches discard data in order to reduce the demand on resources while other strategies avoid the holistic approach and focus on smaller, local subsets of the data, such as in kernel convolution, or Getis & Franklin’s local point pattern analysis. In the current approach, only one point is evaluated by a model at a time and only the chosen number of near neighbours need to be considered, which means very large data-sets can be processed without requiring exorbitant computing resources.

Here, we take the list of point coordinates and for each point determine the Euclidean distances to the specified number of near neighbours. A unique identifier for each point in the set is also assigned. The number of near neighbours considered here reflects the size of the clusters which can be distinguished by the model. For clusters with more points than can be ‘seen’ by the model, their constituent points will be less likely to be classified as clusters, especially those located within the core of the cluster as they will be seen to be within a non-clustered environment based on the context given by the restricted assessment of near neighbours. There is no penalty to over-estimating the number of neighbours to assess as the model will still be expected to see the step-change in input values for clusters with very few points, as the sequence extends to points found outside of the cluster. For the data preparation stage, the distance values for each point are saved to a memory-mapped file for use by subsequent stages. In addition, the unique identifier for each nearby point is also recorded. This is the most resource-heavy aspect of the method as it can require large amounts of storage space if many images are to be analysed, e.g. the distances for 1 million points to their nearest 1,000 neighbours (stored as 64 bit floating point values) would require some 8 gigabytes of storage space.

For each point, the difference between consecutive distances is calculated and then normalized. These normalized distance-difference values are the input used in training of new models or for classification by existing models. The list of distances could itself be used as input (after normalization) to train a model, however although this was as accurate as compared to models trained using the normalized distance-difference values, these models were less stable in the 10-fold crossvalidation testing.

### Model specification and training

Keras, an open-source Python neural-network framework for Python, was used to build and train a neural network model using the distance-difference values as training data and the ‘ground truth’ values to assess performance of the model and produce validation statistics. The distance values from all the simulated data-sets were calculated, as per the data preparation stage, above. Each was loaded alongside the matching ground-truth classification labels. Two thousand distance values for points from each classification label were randomly selected and added to a pool of points for that classification label. If the requisite number of points were not available for a particular label then as many points as possible were sent to the pool. This was repeated for all files in the simulated data-set to yield a final pool of training points evenly sourced from across all the original files, i.e. very large or dense data-sets were not disproportionately represented within the training pool. From each classification label’s training pool a subset of points were randomly selected, pooled, and shuffled (along with their matching classification labels) to create the final training data-sets. A second subset of different randomly selected values was used to created the validation data-set. The models described here were trained on 500,000 points with an even mix of clustered and non-clustered values and validated against 100,000 different values, also with an even mix of cluster type. The training and evaluation data were exported for subsequent use by the model training stage of the workflow.

For the model training stage, a model was assembled from Keras layers and supplied with the training and validation data. Keras training was performed on either a CPU (or a GPU if available) using the TensorFlow back-end(version 1.8.0, (Martín Abadi et al. 2015)). Models were trained over 10 to 100 epochs with a batch size of 32. The ADAM optimizer (Kingma and Ba 2014) was used with an initial model learning rate of 0.001 which reduced with each new epoch and loss determined by binary crossentropy. The model produces a score (from 0 to 1) representing the likelihood that the set of input values were derived from a point situated within a cluster. The score in converted into a label according to a threshold: a score equal or above 0.5 was given the label ‘1’ indicating a clustered point; a score below 0.5 was assigned the label ‘0’ for non-clustered. The model-assigned labels were compared to the ground-truth labels for the simulated data and used to calculate the performance of the model. After training the model was exported to disk along with training and validation performance tracking across the epochs. The performance of the final model was evaluated on the validation data to determine scores for precision, recall, and F_1_ (a combined precision and recall score) and to generate confusion matrices. The model could then also be used in a ten-fold cross-validation assessment to check the performance stability over multiple subsets of the validation data.

### Evaluation of novel data with trained models

Novel data were prepared as for the training data and supplied to a trained model. The model’s scores and classification labels for the data were appended to the original data table and saved to disk. The classification labels were then used to segment the points into like-clusters.

### Post-evaluation cluster segmentation and shape fitting

Individual clusters were extracted from the set of each points’ x-y coordinates, classification label, near-neighbour distances, and near-neighbours’ unique identifiers. Clustered points were grouped into similar clusters by considering each clustered point in turn, assigning that point a unique ‘Cluster ID’ and then considering all the nearby points in order of their proximity to the original point. Nearby points which were also labelled as ‘clustered’ were assigned the same Cluster ID and this process continued until the first nearby point was reached which was labelled as ‘non-clustered’. If any of the nearby-clustered points have already been assigned a Cluster ID then all points adopt that Cluster ID (if there are several other Cluster IDs then the lowest Cluster ID is adopted by those points. All other points with the competing Cluster IDs are also reassigned in this manner). Otherwise all points keep the original Cluster ID and the process is repeated with another clustered point until all clustered points have been assigned a Cluster ID. This method therefore utilises all the labelling information from the model: clustered points are used to initiate the growth of a new cluster and non-clustered points used to halt the process. However, this approach will only succeed while there are non-clustered points present to indicate when a cluster should stop acquiring new points. In cases where a large majority of the points are clustered there are two approaches that can be used to segment clusters which are separated by empty space rather than non-clustered points. First, in cases where all the nearest points (as many as were measured in the pre-processing step) were labelled as ‘clustered’ by the model, an additional check can be triggered, which considers the spatial arrangement of these points. Here the mean distance to the nearest three points is taken and compared against the mean as more consecutive nearby points are included. If the mean distance increases by more than 5 times the standard deviation of the mean then the cluster is ‘split’ at the previous point and all earlier points assigned the same Cluster ID. Once all clustered points have been assigned a Cluster ID, each Cluster ID is checked to ensure it contains at least three points. Points are relabelled as non-clustered if their Cluster ID fails this test. Next, each set of like-clustered points (sharing the same Cluster ID) was used to fit an encapsulating cluster shape.

A set of points has no intrinsic area but a polygon can be constructed which contains the set of points and approximates the intuitive shape of the spatial distribution of those points, including spurs and (with additional polygons) holes and other features. Such a polygon can be created by several methods including bounding box, convex- and alpha-hulls, however these methods can not always produce a shape which satisfactorily describes the distribution of the points. The method used here involves dilating each point in the set by a certain radius, then taking the union of all the overlapping discs from each dilated point to form the cluster polygon. The selection of the dilation radius is an important parameter in this process, for if the dilation radius is too small then the shapes formed from points may not overlap and the ‘cluster’ fragments into disconnected polygons. If the dilation radius is too large then the resultant shape contains all the points but is a poor representation of the spatial distribution of those points. A suitable radius can be estimated from the ratio of the bounding-box area of the set of points to the number of points in the set. In this way, the dilation radius adapts to the area over which the points are spread and the number of points which need to be enclosed. To reduce the extent to which the resultant shape’s boundary falls beyond the ‘outer’ points, the cluster polygon can also be eroded by a fraction of the dilation radius to yield a final shape which contains all the prescribed points in a relatively free conformation.

This method may generate multiple polygons if, for example, in the case where two clusters were close enough to be considered as one by the segmentation process but are separated by a distance greater than the dilation radius used to create the cluster outline. In these cases, polygons containing more than a specified number of points (again here we used three or more points) can be considered as separate clusters and their constituent points given separate new unique Cluster IDs. Polygons containing fewer points are not considered to be clusters and their points can be stripped of the Cluster ID and their cluster label re-assigned to ‘nonclustered’.

This method may also generate polygons containing holes; these holes can either be preserved or removed to close the hole. When closing holes it is possible to specify a relative size threshold below which holes are closed and above which the holes are preserved in order to better represent, for example, ring-shaped clusters. Here we removed holes which were smaller than half of the dilation radius used to construct the cluster shape.

The final cluster shape can be assessed for various shape descriptors such as the cluster area, perimeter length, and circularity ratio. As the number of points in the cluster is known further metrics such as the density of points within the cluster can be determined. As the entire original image is processed, thousands of clusters can be assessed and slight differences in cluster distributions relative to particular areas of the cell may become apparent and these can be further examined using targeted regions of interest.

### Extraction of cluster information from regions of interest

Regions of interest in each input image were obtained by a freehand selection tool written in MATLAB (Mathworks). Following region selection, the data produced by the model assessment and cluster segmentation stages can be loaded and information on clustering within each ROI extracted and saved using a Python script.

### Cell isolation & staining

Primary human T-cells were isolated using a pan T cell selection kit (130-096-535, Miltenyi Biotec). Untouched T cells (‘naive cells’) were isolated from human blood and then used directly for imaging assays. T blasts (‘pre-stimulated cells’) were generated in parallel by incubating untouched T cells with anti-CD3 (RnD Systems, Clone OKT-3 at 2 μg/ml) and anti-CD28 (RnD Systems, clone CD28.2 at 5 μg/ml) coated flasks in complete medium (RPMI with 10% fetal bovine serum, glutamine, and penicillin/streptomycin) for two days. Cells were then washed and cultured in complete medium for an additional 5 days in the presence of 20 ng/mL IL-2 (Proleukin). Purity was then assessed by FACS analysis.

Glass-bottomed chamber slides (#1.5 glass, ibidi μSlides) were coated with a mixture of recombinant human ICAM-1-Fc (RnD Systems at 3 μg/ml) and anti-CD3 antibody (2 μg/ml) overnight at 4°C. Cells were added to wells at a density of 25,000 cells/cm^2^ for 4 minutes then gently rinsed with warm (equilibrated to 37°C) HBBS to remove non-adhered cells then fixed. Fixation was by the pH-shift method at room temperature: cells were first incubated for 5 minutes in pH 6.8 Fixation Buffer (3% (w/v) para-formaldehyde (PFA), 80 mM PIPES, 2mM MgCl_2_, 5 mM EGTA, in water) followed by 10 minutes in pH 11 Fixation Buffer (3% (w/v) PFA, 100 mM Borax, in water). Fixed cells were washed three times with PBS, permeabilized with Triton X-100 (0.1% in PBS) for 5 minutes at 4°C and rinsed again. Auto-fluorescence was quenched by incubating the sample in NaBh_4_ (1 mg/ml in water) for 15 minutes followed by rinsing three times with PBS. The fixed, quenched samples was blocked with 5% (w/v) BSA/PBS for an hour. The sample was then incubated with primary antibody, either rabbit polyclonal anti-Csk (Santa Cruz sc-286) at 1:300 or rabbit polyclonal anti-PAG (Abcam ab14989) at 1:500 overnight at 4°C and washed three times for 5 minutes with PBS. The sample was then incubated with secondary antibody labelled with anti-mouse Alexa Fluor 647 (ThermoFisher Scientific) for 1 hour at room temperature followed by three 5 minute PBS washes. The sample was then used immediately for dSTORM imaging.

For the non-stimulatory condition (referred to as ‘glass’), cells were fixed, washed, and stained as a single-cell suspension. Fixed, stained cells were settled onto untreated glass chamber slides prior to imaging.

### dSTORM image acquisition

Fixed and stained samples were prepared for imaging by replacing the final PBS wash with a volume of STORM imaging buffer (50 mM Tris-HCI (pH 8.0), 10 mM NaCl, 0.56M glucose, 0.8 mg/ml glucose oxidase (Sigma G2133), 42.5 μg/ml bovine catalase (Sigma C40), 10 mM cysteamine (Sigma 30070). The dSTORM image sequences were acquired on a Nikon N-STORM system in a TIRF configuration using a 100× 1.49 NA CFI Apochromat TIRF objective for a pixel size of 160 nm. Samples were illuminated with 647 nm laser light at approximately 2.05 kW/cm2. Images were recorded on an Andor IXON Ultra 897 EMCCD using a centered 256 × 256 pixel region at 20 ms per frame for 30,000 frames and an electron multiplier gain of 200 and pre-amplifier gain profile 3.

### dSTORM image reconstruction

The dSTORM imaging data were processed using ThunderSTORM (Ovesný et al. 2014) and the following parameters: pre-detection wavelet filter (B-spline, scale 2, order 3), initial detection by non-maximum suppression (radius 1, threshold at one standard deviation of the F1 wavelet), and sub-pixel localisation by integrated Gaussian point-spread function (PSF) and maximum likelihood estimator with a fitting radius of 3 pixels. Detected points were then filtered according to the following criteria: an intensity range of 500 - 5000 photons, a sigma range of 50 - 250, and a localisation uncertainty of less than 25 nm. The filtered data-set was then corrected for sample drift using cross-correlation of images from 5 bins at a magnification of 5. Repeated localisations, such as can occur from dye re-blinking, were reduced by merging points which re-appeared within 50 nm and 25 frames of each other, in the manner of (Annibale et al. 2011). For the purposes of interpretation it is assumed that the prevalence of re-blinking dye molecules and the proportion that escape the merging process is independent of the sample staining and therefore the relative changes in the clustering of points between conditions are independent of dye re-blinking.

### Statistical Analyses

For statistical comparisons, all data were analysed using non-parametric Kruskall-Wallis and Dunn’s multiple comparison tests in Graphpad Prism software. A significant difference between conditions was considered to be P < 0.05 for rejecting the null hypothesis.

## Results

### Machine learning models can be trained to accurately identify clustered points

Training data comprised of a set of files generated by the cell-simulator which covered a broad range of different clustering scenarios. These scenarios were compiled from combinations of different clustering parameters, specifically an overall point density of 5, 10, 50, 100, 300, or 500 points per μm^2^, a point density within each cluster of 5, 10, 20, 30, 40, 50, 60, 70, 80, or 90 points per cluster, a proportion of clustering of 5, 10, 20, 30, 40, 50, 60, 70, 80, 90, or 100 percent of points being in clusters, and a maximum distance from the cluster seed (effectively the cluster radius) of 5, 10, 15, 25, 40, 50, 75, or 100 nm. Each clustering parameter combination therefore describes a ‘clustering scenario’, however some combinations yield untenable scenarios. For example: a very high density of clusters with a large radii cannot be accommodated with the additional constraint that clusters should not overlap. Similarly, the density of points within clusters containing a low number points distributed within a large enough radii can become indistinguishable from the density or distribution of points outside of the cluster, or if the density inside clusters is much lower than that outside the clusters can appear as ‘holes’ rather than ‘clusters’. Therefore only clustering scenarios which resulted in 1 to 5 cluster(s) per μm^2^ and with a point-density inside clusters of between 1.5× and 100× that of the non-clustered point-density were considered as viable scenarios for simulation. Of the 5,280 possible combinations of the specified clustering parameters there were 711 viable scenarios; the remaining non-viable scenarios were excluded from simulation. Each viable scenario was used to distribute points within a simulated cell-synapse outline. Completely spatially random data (without any clustered points) were also simulated and included for the training of some models. Training data files were then used as input for the pre-processing stage where the nearest neighbour distances were recorded. For models 07VEJJ the 1^st^ to 100^th^ nearest neighbours were considered. For models 87B144 and XPILJZ the 1^st^ to 1000^th^ nearest neighbours were assessed. These data were then used to construct training data-sets as described earlier. Preparing data-sets for 1000 nearest-neighbours from input sets containing approximately 1 million, half a million, or one hundred thousand points took approximately 84, 23, and 1.6 minute(s), respectively, on a Core i7-4930K personal computer with 32 GB RAM.

Model 07VEJJ was configured with 12 layers in Keras as described earlier and trained on 500,000 input samples comprising an even mix of clustered and nonclustered labels. The model was validated on 100,000 different input samples with the same even mix of classification labels. This model demonstrated an accuracy of 92.36% on the training data and 92.44% on validation data, with an F1 score of 0.9243 for the validation data (precision 0.9245 and recall 0.9243). Ten-fold cross validation of the model resulted in an accuracy score of 91.81% ± 0.27%. Other models used in this study achieved similar performance: Model 87B144 (12 layers using 1000 near-neighbours) showed an accuracy of 94.01% on the training data and 93.99% on validation data with an F1 score of 0.9398 for the validation data (precision 0.9420 and recall 0.9399). Model XPILJZ (4 layers using 100 near-neighbours) showed an accuracy of 91.86% on the training data and 91.99% on validation data with an F1 score of 0.9199 for the validation data (precision 0.9204 and recall 0.9199). Processing time for these models was linear; Model 07VEJJ took 6.6 minutes to evaluate 1 million input points (each with 100 nearest-neighbour values) and Model 87B144 took 50.7 minutes to process the same data-set (using 1000 nearest neighbour values) using an NVIDIA GeForce GTX 750 Ti GPU with 2GB RAM. Subsequent performance times depend on the number and complexity of the clusters identified after points are classified by a model.

**Figure 3.**
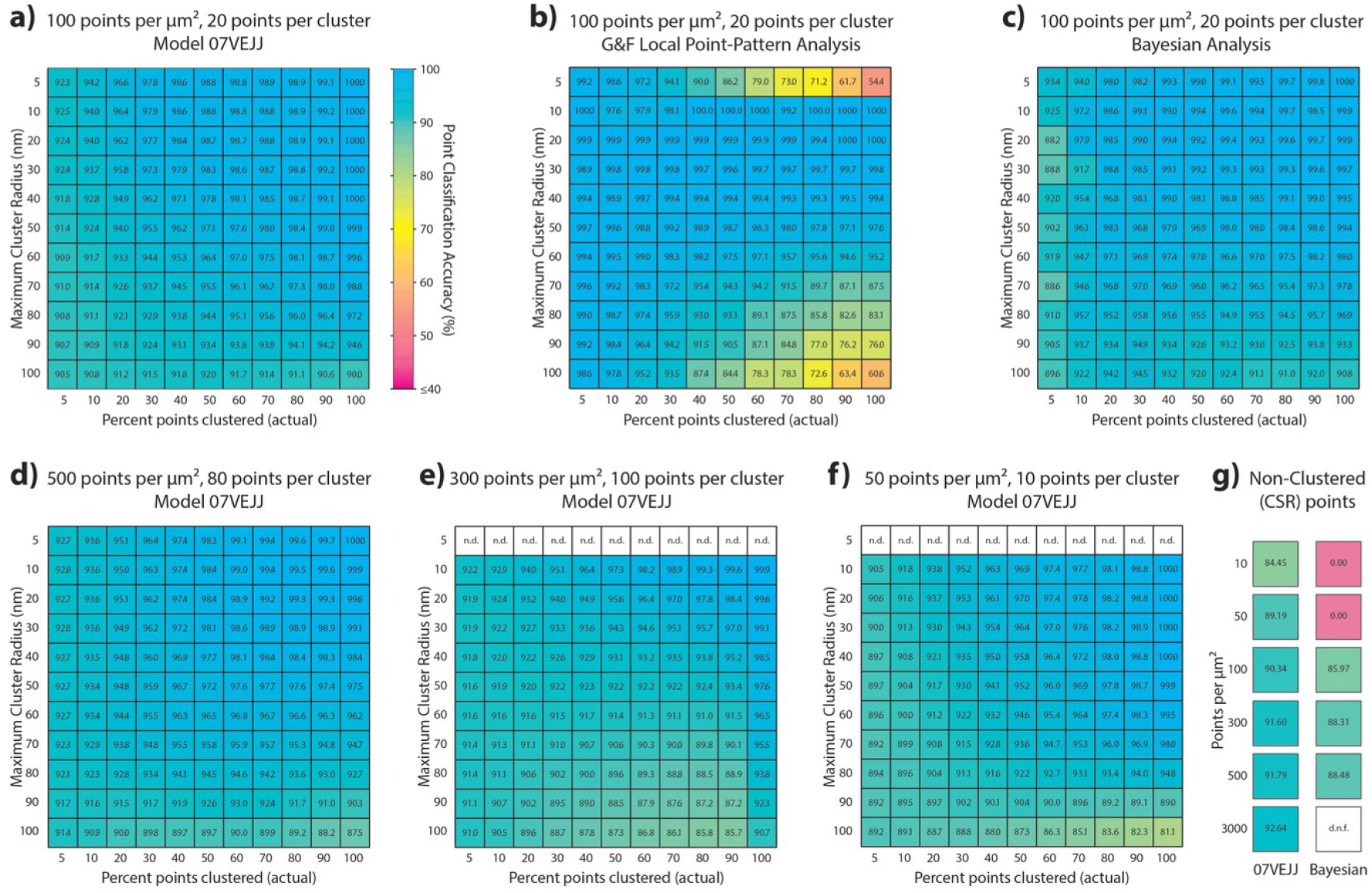
The classification accuracy of validation data assessed with (a) Model 07VEJJ or (b) Getis & Franklin’s Local Point Pattern Analysis or (c) Bayesian Cluster Analysis on 3 × 3 μm centred regions extracted from the data-sets analysed by (a) and (b). Model 07VEJJ shows comparable performance at very high point-densities (d), when the number of points per cluster matches the limit of the models near-neighbour distances ‘reach’ (e, n.d. = not determined), and very low point-densities (f). (g) Classification accuracy for completely spatially random data at different point densities analysed by Model 07VEJJ or the Bayesian method; d.n.f = did not finish.

The model was then used to evaluate novel simulated data, including images only containing spatially random (non-clustered) points (Fig. 3a, g). These data featured the same range of clustering scenarios as that which was used to derive the training and validation data-sets. Model 07VEJJ showed greater than 90% accuracy in CSR data where the overall point density was 100 points per μm^2^ or greater. For lower-density data the accuracy decreased to 89.19% in images with 50 points per μm^2^ and 84.45 % for those with 10 points per μm^2^. In simulated data-sets with clustering present, 07VEJJ was able to correctly identify labels for over 90% of points in the majority of clustering scenarios (Fig. 3d, f). Accuracy was lower when clusters became very large or when only a small proportion of the points were in clusters. In the latter scenario the model was often found assigning clustered labels to nonclustered points at a similar rate to the incorrect identification of clustered points in CSR data. When presented with data outside of the original training data clustering scenarios, for example clusters containing 100 points (also note that Model 07VEJJ can only ‘see’ as far as the 100 nearest neighbours) the model was still able to perform the correct classification in approximately 90% of the cases (Fig. 3e).

The simulated data-sets were also used to evaluate the performance of different cluster analysis methods. A stalwart method is that of Getis & Franklin: their local point-pattern analysis (G&F LPPA) is a spatially restricted version of the original Ripley’s *K* Function and Besag’s *H* Function, in that it only considers the density of points at a single fixed radius. Each point acquires a value indicating if the local point density (within the fixed radius) is greater or less than the expected point density; this value is effectively a measure of the point’s clustering behaviour at the selected spatial scale. This value can then be subjected to a threshold to classify points as clustered (above threshold) or non-clustered (below threshold). However the specification of the radius and threshold are very much dependent on the underlying data and are sensitive to multiple scales of clustering and inhomogeneities (for example a very large dense cluster of points somewhere in the data can skew the assessment of the ‘expected point density’ considerably). As the simulated data-sets contain only a single type of cluster which is evenly distributed through the ‘cell area’ a G&F LPPA analysis should not be too perturbed. The analysis was favoured by using a radius which was matched to the expected cluster radius for each specific data-set. A threshold was chosen by first analysing five data-sets of the equivalent number of points as the original, but with x-y coordinates that were spatially randomised within the cell-shape. The 99.5 percentile value for all L(r) values from all five randomised images was used as the threshold value for the original data-set. When applied to all the simulated data-sets, the G&F LPPA performed reasonably well but also performed worst on images with a high proportion of points in either very small or very large clusters. Overall the performance of this method, even with the parameters individually optimised for each image, was worse than Model 07VEJJ (Fig. 3b).

Another cluster analysis method utilizes Bayesian model-based inference to select the best radius and threshold pairing from among thousands of possible radius-threshold combinations. Each combination is examined as with G&F local point pattern analysis and then scored according to how well it matches a Bayesian generative model with points distributed in circular clusters according to a Dirichlet process (Rubin-Delanchy et al. 2015; Griffié et al. 2016). This method can be quite computationally intensive for data-sets with more than approximately 25,000 points and to this end, a 3 × 3 μm region from the centre of each simulated data-set was extracted for analysis. When applied to these subsets of the simulated data, the Bayesian approach performed as well or slightly better than Model 07VEJJ, which was assessed on its performance across the full data-set (Fig. 3c). However for data simulated with a low overall point-density of 10 or 50 points per μm^2^, the Bayesian method failed to correctly classify points in all many cluster scenarios, instead labelling all points as clustered in all scenarios. For this reason, the performance of the Bayesian analysis appears to improve as the percentage of points in clustering increases. When comparing spatially random points (i.e. no clustering) of different points densities, Model 07VEJJ showed slightly better performance than the Bayesian for those conditions in which the Bayesian was able to operate (Fig. 3g). As with the clustered data, all points from conditions with low point density were mislabelled as clustered. For high-density data (3000 points per μm^2^) the Bayesian method was unable to finish processing any of the 3 × 3 μm region subsets.

### Clustering of Csk and PAG proteins in T cells

Csk is a protein tyrosine kinase and a well established negative regulator of T cell receptor signalling through its inactivation of membrane-associated Src kinases such as Lck. As a cytosolic protein, Csk is through to be regulated through its recruitment to the transmembrane adapter protein PAG. In non-activated T cells PAG is predominantly phosphorylated (Brdicuka et al. 2000; Davidson et al. 2003) which facilitates Csk binding. Membrane-proximal Csk is therefore well positioned to exert its inhibitory effects upon membrane-associated Src kinases. Upon TCR stimulation Csk unbinds from PAG, possibly as a result of changes in the phosphorylation state of PAG (Reginald et al. 2015; Torgersen et al. 2001), leading to a more permissive environment for Src kinase activation.

Primary human T cells demonstrated changes in the clustering of Csk and PAG at the plasma membrane (Fig. 4). These changes were dependant on both the status of the T cells (naive or pre-stimulated) and their activation status (non-activated or activated on anti-CD3+ICAM-1 coated glass).

### Csk Clustering

Naive cells showed a change in Csk clustering with surface-activation through an increase in the number of clusters detected by Model 87B144 from 4.85 clusters per per μm^2^ (median, IQR 3.15 - 37.38 clusters per μm^2^) for non-stimulated cells to 8.80 clusters per μm^2^ (median, IQR 6.75 - 10.75 clusters per μm^2^) for cells stimulated on anti-CD3+ICAM-1 coated glass, P < 0.0001. These data were not significantly different to that reported from a Bayesian analysis (P > 0.9999 for both stimulation conditions).

Pre-stimulated cells showed a change in Csk clustering after cell activation as an increase in the number of clusters detected by Model 87B144 from 6.77 clusters per per μm^2^ (median, IQR 5.13 - 8.37 clusters per μm^2^) for non-stimulated cells to 14.37 clusters per μm^2^ (median, IQR 11.03 - 18.57 clusters per μm^2^) for cells stimulated on anti-CD3+ICAM-1 coated glass, P = 0.0009. Compared to the same data analysed with the Bayesian method, there was no significant different for the stimulatory condition (P> 0.9999), however the Bayesian analysis reported fewer clusters in the non-stimulatory condition at 3.78 clusters per per μm^2^ (median, IQR 2.11 - 6.42 clusters per per μm^2^), P = 0.0006.

Naive cells also showed an increase in the percentage of points clustering from 53.21 % (median, IQR 48.33 - 59.69 %) in the non-stimulated condition to 61.15 % (median, IQR 53.14 - 64.39 %) for the stimulated condition, P = 0.0023. The Bayesian analysis identified the same trend but returned significantly lower values than CAML with 46.86 % (median, IQR 41.86 - 52.43%) points clustered in the nonstimulatory condition (P < 0.0001 compared to CAML) increasing to 52.82 % (median, IQR 44.35 - 59.92%) points clustered in the stimulatory condition (P < 0.0001 compared to CAML), P < 0.0001 (comparing stimulatory condition).

Pre-stimulated cells did not significantly change the percentage of points clustering between stimulatory conditions with 54.01 % (median, IQR 49.82 - 59.47 %) of points clustered in the non-stimulated condition and 59.12 % (median, IQR 54.82 - 67.92 %) points clustered for the stimulated condition, P = 0.2012. This was confirmed in the Bayesian analysis although, as for the naive cells, there was a reduced percentage of Csk in clusters reported by this method, with 55.30 % (median, IQR 45.75 - 63.90 %) points clustered in the non-stimulatory condition and 53.51 % (median, IQR 44.32 - 66.40 %) points clustered in the stimulatory condition, P > 0.9999 (comparing stimulatory conditions). There was no significant difference between the two analysis methods for the non-stimulatory condition (P > 0.9999) and a slight difference for the stimulatory condition (P = 0.05).

When considering the number of points per cluster, naive cells showed an increase from 9 (median, IQR 6 - 14) points per cluster in the non-stimulatory condition to 11 (median, IQR 7 - 18) points per cluster for the stimulatory condition (P < 0.0001). The Bayesian analysis reported significantly more points per cluster for the same data (P < 0.0001 comparing analysis methods within either stimulation condition) however the method returns a mean value for each ROI whereas CAML returns a value for each cluster; for the non-stimulatory condition the Bayesian method reported 14.08 (median of ROI means, IQR 11.44 - 17.47) points per cluster increasing to 16.87 (median of ROI means, IQR 14.35 - 20.84) points per cluster after stimulation (P = 0.0046).

Pre-stimulated cells also increased the number of points per cluster with 12 (median, IQR 7 - 22) points per cluster in the non-stimulatory condition rising slightly to 14 (median, IQR 8 - 27) points per cluster for the stimulatory condition (P < 0.0001). The number of points per cluster was also higher in pre-stimulated cells compared to naive cells regardless of the activation condition of the cells (P < 0.0001 in either case). This trend was reflected in the Bayesian analysis of the same data although, as before, the mean ROI values were higher; 15.16 (median of ROI means, IQR 10.35 - 19.92) points per cluster increasing to 24.00 (median of ROI means, IQR 20.13 - 28.99) points per cluster in the stimulatory condition (P < 0.0001). With the alternative analysis method, there were no significant differences between the naive and pre-stimulated cells on the non-stimulatory condition (P > 0.9999) but still significantly more points per clustering in the pre-stimulated cells than the naive cells for the stimulatory condition (P < 0.0001).

The area of Csk clusters in naive cells decreased slightly from 2322 (median, IQR 1310 - 4522) nm2 when non-stimulated to 2144 (median, IQR 1261 - 4522) nm2 after stimulation (P < 0.0001). This trend was exacerbated for pre-stimulated cells where clusters shrank from 3093 (median, IQR 1632 - 6154) nm2 in non-stimulated cells to 1920 (median, IQR 1030 - 3655) nm2 with stimulation (P < 0.0001). Comparing between cell status, the pre-stimulated cells increased their Csk cluster area compared to naive cells for non-stimulatory conditions (P < 0.0001) but Csk clusters in per-stimulated cells became much smaller upon anti-CD3+ICAM-1 stimulation (P < 0.0001).

Combining the measurements of cluster size and the number of points per cluster, the density of points for each cluster can be determined. For naive T cells, the density within clusters increased from 0.003921 (median, IQR 0.002206 - 0.006558) points per nm2 in non-stimulatory conditions to 0.004591 (median, IQR 0.002924 - 0.007082) points per nm2 with stimulation (P < 0.0001). For prestimulated T cells, the density within clusters also increased from 0.003939 (median, IQR 0.002517 - 0.006243) points per nm2 in non-stimulatory conditions to 0.004657 (median, IQR 0.002846 - 0.007249) points per nm2 with stimulation (P < 0.0001). There was no difference in the density of points within clusters between cell status for the non-stimulatory condition (P > 0.9999), however the density was greater for pre-stimulated cells that for naive, after anti-CD3-ICAM-1 stimulation (P < 0.0001).

### PAG Clustering

Naive cells showed a change in PAG clustering with surface-activation through an increase in the number of clusters from 10.19 clusters per per μm^2^ (median, IQR 5.03 - 16.07) clusters per μm^2^) for non-stimulated cells to 13.89 clusters per μm^2^ (median, IQR 7.07 - 21.32 clusters per μm^2^) for cells stimulated on anti-CD3+ICAM-1 coated glass, P = 0.0002. Pre-stimulated cells did not show a change in PAG clustering between cell activation conditions with 1.20 clusters per per μm^2^ (median, IQR 0.53 - 1.59 clusters per μm^2^) for non-stimulated cells and 0.99 clusters per μm^2^ (median, IQR 0.78 - 1.10 clusters per μm^2^) for cells stimulated on anti-CD3+ICAM-1 coated glass, P > 0.9999. These data also show a considerable decrease in PAG clustering, in both non-activated and activated conditions, for pre-stimulated cells compared to naive cells (P < 0.0001).

Naive cells did not show a change in the percentage of points clustering between non-stimulated and stimulated conditions, with 50.45 % (median, IQR 47.39 - 54.45 %) points clustered in the non-stimulated condition and 54.57 % (median, IQR 50.36 - 59.09 %) for the stimulated condition, P = 0.1333. Pre-stimulated cells also did not significantly change the percentage of points clustering between stimulatory conditions with 64.65 % (median, IQR 59.77 - 68.77 %) of points clustered in the non-stimulated condition and 63.99 % (median, IQR 60.24 - 66.94 %) points clustered for the stimulated condition, P > 0.9999. Comparing between cell status, naive cells had fewer percentage of the Csk points in clusters that pre-stimulated cells in either stimulation scenario (P < 0.0001).

When considering the number of PAG points per cluster, naive cells showed no apparent change with 10 (median, IQR 6 - 18) points per cluster in the nonstimulatory condition to 10 (median, IQR 6 - 18) points per cluster for the stimulatory condition. However, the distributions of these two conditions was statistically different (P > 0.0001). Pre-stimulated cells, although presenting far fewer clusters overall, showed a decrease between the non-stimulatory condition, with 14 (median IQR 7 - 32) points per cluster compared to the stimulatory condition with 11 (median, IQR 6 - 20) points per cluster (P < 0.0001). In comparing between the two types of cells there were more points per cluster in pre-stimulated cells than in naive for the non-activating conditions (P < 0.0001). There was no difference in the number of points per cluster for the activating conditions (P = 0.0509) between naive or pre-stimulated cells.

The area of PAG clusters in naive cells decreased slightly from 1551 (median, IQR 840.9 - 3159) nm2 for non-stimulated cells to 1359 (median, IQR 766 - 2522) nm2 after stimulation (P < 0.0001). In pre-stimulated cells, the cluster areas remained similar with 19504 (median, IQR 8597 - 41690) nm2 in non-stimulated cells and 15669 (median, IQR 8054 - 30041) nm2 in stimulated cells (P > 0.9999). Comparing between cell types, the pre-stimulated cells had PAG clusters with considerably greater area compared to naive cells for both non-stimulatory and stimulatory conditions (P < 0.0001 in both cases).

For naive T cells, the density within clusters increased from 0.006828 (median, IQR 0.004383 - 0.01034) points per nm2 in non-stimulatory conditions to 0.008138 (median, IQR 0.005635 - 0.01165) points per nm2 with stimulation (P < 0.0001). For pre-stimulated T cells, the density within clusters was not different between stimulation conditions, with 0.0008665 (median, IQR 0.000568 - 0.00132) points per nm2 in non-stimulatory conditions and 0.000782 (median, IQR 0.000509 - 0.001214) points per nm2 with stimulation (P > 0.9999). Concerning the differences between cell types, there was again a considerable decrease in point-density within clusters for pre-stimulated cells compared to naive (P < 0.0001) for both stimulation conditions. This is expected considering the other observations of considerably larger clusters in pre-stimulated cells without much change in the number of points within each cluster.

**Figure 4.**
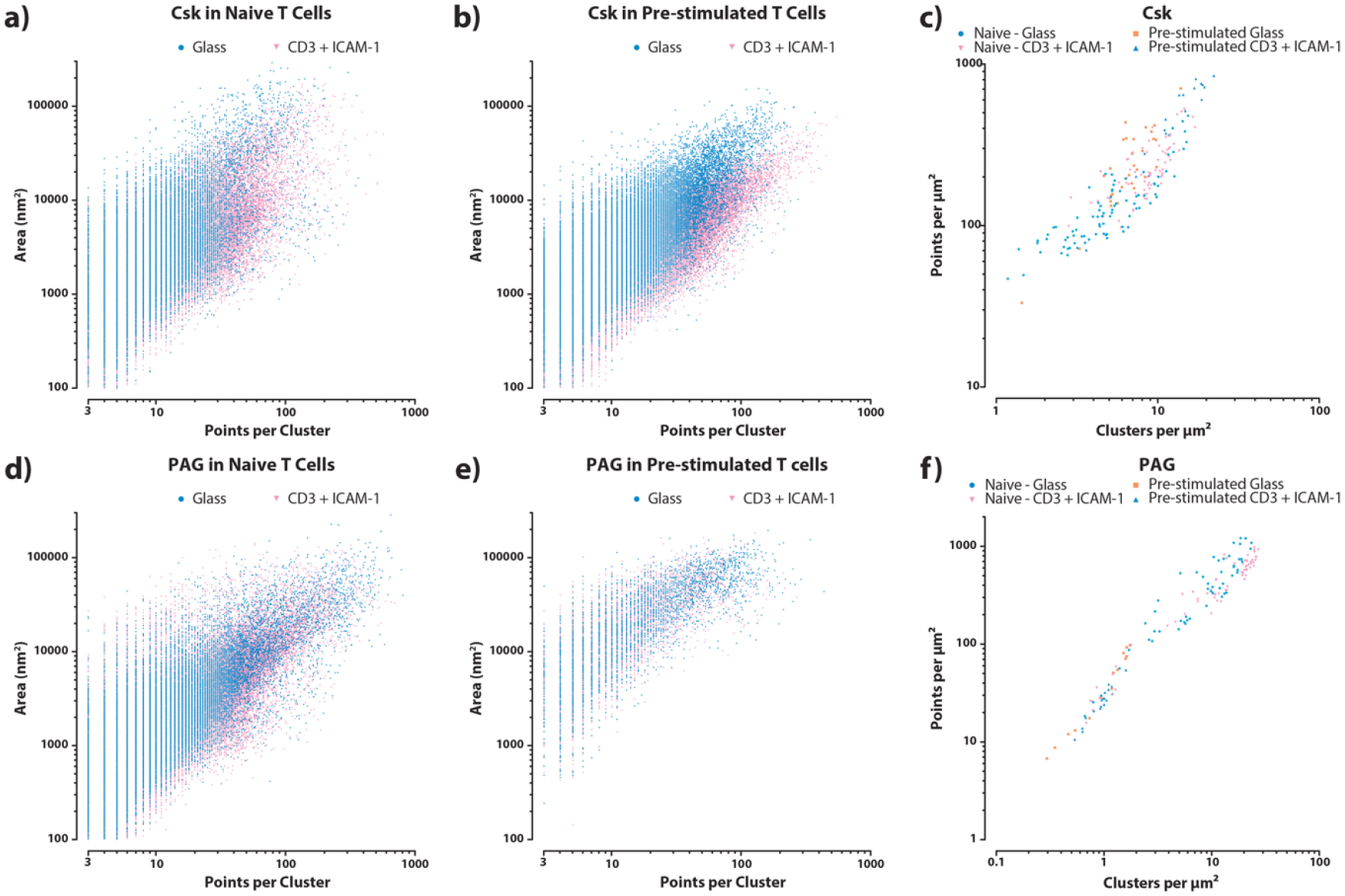
Cluster metrics from CAML (model 87B144) for dSTORM images of Csk (top row) and PAG (bottom row). a,d) Area of, and points contained by, individual clusters detected in naive T cells which were either unstimulated (blue circles) or stimulated on anti-CD3 + ICAM-1 surfaces (pink triangles). b, e) Area of, and points contained by, individual clusters detected in Pre-stimulated T cells which were either unstimulated (blue circles) or stimulated on anti-CD3 + ICAM-1 surfaces (pink triangles). c, f) Overall density of clusters and points within whole-cell regions of interest for unstimulated naive T cells (blue circles), unstimulated pre-stimulated T cells (orange squares), stimulated naive T cells (pink down-triangles), or stimulated pre-stimulated T cells (blue up-triangles).

## Discussion

In this study we used machine learning networks to classify points from SMLM datasets as either clustered or non-clustered, based solely upon a sequence of values derived from each point’s nearest-neighbour distances. The network is able to identify features within such a sequence which allow it to classify points with a high degree of confidence. In certain cases of low-density input images or a low percentage of clustering, the model tended to overestimate the clustering of points. For images with a low percentage of clustering, i.e. a high proportion of spatially randomly distributed points, the excess clustering can be attributable to the model finding groups of points which happen, by chance, to be spatially closer than their further neighbours, For low-density images, in particular those simulated images with the few points contained within very large clusters, then the overall clustering pattern breaks down and these large-but-sparse clusters become difficult to discern from spatially randomised data.

We have also shown how data-sets which have been annotated by our model can then be further processed to use the new information on a point’s clustering status to partition points into separate clusters. Once partitioned, cluster outlines can be drawn and additional information on clustering extracted. This process can be performed on the entire available data-set, without needing to reduce the data or acquire expensive computational resources. These per-cluster data allow for a deeper interrogation of the clustering patterns within the original image compared to global reporters of clustering, such as Ripley’s *K* Function. This is of particular relevance to data from biological specimens which, without reduction of the original data to small regions of interest, is rarely homogenous in its overall distribution of either points or clusters.

Our approach has several benefits, in particular that it is comparatively fast compared to other available methods and requires minimal parameter input. While the processing turnaround time is not a linear function of the number of input points, it is still possible to process heavily populated data-sets within a comfortable time-frame on a modest desktop computer. Furthermore the only analysis parameter which bears any major consequences on the output is the number of nearest neighbours from which distances are measured. This decision is easily informed from a cursory examination of the input data and an overestimation of this parameter does not have a detrimental effect on the outcome. Underestimation results in the exclusion of clusters containing more points than the model can ‘see’; for samples containing fiducial registration beads (for drift and aberration correction or channel alignment) this could be employed to remove beads, which often present as very large, very densely populated clusters within the reconstructed image.

Although the models described here were trained on simulated clusters with hard-edges and circular shapes, which may be somewhat removed from those seen in biological samples imaged by common SMLM methods, they are nevertheless able to find clusters of abstract and arbitrary shapes, including highly elongated clusters from fibrous structures. Our models’ assessment of clustering is also not affected by discontinuities in the distribution of points or the type of clustering within the field of view and it is not limited or specialised in finding clusters of a particular shape. The method as described here is flexible enough to allow an end user to quickly assemble a new model with training data generated to match a particular of clustering outcome, while also allowing rapid re-use of trained models to classify and annotate data for the quantification of clustering in ‘real’ data.

Limitations of this are approach are, in the main, that any model cannot see clusters which contain many more points than the number of near-neighbour distances which are supplied as input. Points within such large clusters will likely be incorrectly labelled as non-clustered. This is particularly the case for points situated within the core of the cluster where their near-neighbour distances cannot incorporate any clustering features for a model to recognise, or at least none of the scale of the main cluster leading to the segmentation of several smaller, tightly aggregated clusters within the main cluster. Points on the edge of a very large cluster may still be recognised correctly by models with limited near-neighbour ranges, as the near-neighbour points will include those from the large cluster and some points from outside of the cluster. Together, these two model evaluation artefacts—ring shaped clusters and tight clusters-of-clusters—can be useful signs, when constructing a new model, to recognise if the number of neighbours needs to be increased for a set of data.

The point patterns within SMLM data are highly amenable to annotation by machine learning models, as shown here to extract spatial clustering patterns from both simulated and biologically derived data. Future directions for such work might include incorporating more features from the input data, such as relative angles to near neighbours or including the localisation precision of nearby points, in order to more robustly determine the extent of local point clustering. This will certainly require more sophisticated training data, which are simulated to incorporate these additional features. Here we have used a simple binary classifier—clustered or non-clustered—yet it would be possible to expand the number of classifiers and train a model learn an abstract representation of different types of clusters, for example to label points as situated within a ‘circular cluster’, a ‘fibrous cluster’, a ‘ring-shaped cluster’, or any cluster with particular attributes that can be generated and labelled within the training corpus. Further improvements could be made if the model were to return a vector indicating not just if a point originated from a cluster but also which of the nearby points were likely to come from the same cluster. There is also the potential to expand the method into multidimensional data, such as 3D spatial clustering, dynamic clustering in live data, or co-clustering between points from different imaging channels. The software presented here describes a complementary approach to the existing methods for the cluster analysis of SMLM data. It is presented so as to be easily accessible to non-experienced users while providing flexibility to enable different and highly customised configurations if required.

## Code Availability

The Python scripts and trained models that are described here, as well as a user guide, are available online at https://gitlab.com/quokka79/caml.

